# Single-Cell and Spatial Profiling of Tumor Microenvironment Heterogeneity in Human Osteosarcomas

**DOI:** 10.1101/2025.06.27.661729

**Authors:** Xuejing zheng, Jin Yuan, Peng Jin, Xuejun Wang, Runlei Xu, Helin Feng, Liangliang Wu, Yongjie Xie, Xinxin Zhang, Wence Wu

## Abstract

Osteosarcoma, a highly heterogeneous malignant bone tumor, exhibits substantial interpatient and intratumoral heterogeneity. Utilizing integrated single-cell RNA sequencing and spatial transcriptomics, we uncovered distinct tumor microenvironment (TME) transcriptional profiles across patients, highlighting profound interpatient heterogeneity. Intratumorally, malignant cells primarily followed two differentiation trajectories, converging into osteoblastic and chondroblastic functional states, and hypoxia may be the influencer; notably, within individual patients, tumor cells exhibited greater transcriptional similarity driven by specific transcription factors, despite these divergent states. Spatial analysis revealed patient-specific TME cellular co-localization patterns, alongside conserved spatial relationships: vascular components (endothelial cells, pericytes) demonstrated strong co-localization, while immune cells (T cells, myeloid cells) clustered within shared regions. Crucially, these functional states occupied distinct microniches: osteoblastic regions were enriched with osteoclasts, vascular components, and immune cells, whereas chondroblastic regions displayed the opposite composition and were preferentially located in hypoxic, vascular-poor niches, exhibiting significant enrichment of hypoxia-related signaling pathways. Furthermore, our data suggest osteosarcoma cells may activate fibroblasts via the SPP1 signaling pathway, indicating fibroblasts act as key intermediaries in tumor-directed TME modulation. This study comprehensively delineates the intricate landscape of osteosarcoma heterogeneity, defining distinct functional cellular states, their spatially organized TME niches, and a potential SPP1-mediated tumor-fibroblast regulatory axis. This comprehensive analysis elucidates the intricate interpatient and intratumoral heterogeneity in osteosarcoma, revealing functional and spatial organization within the tumor and its microenvironment.

## Introduction

Osteosarcoma, the most common primary malignant bone tumor, originates from primitive mesenchymal cells and is histologically characterized by malignant spindle cells producing osteoid or immature bone ^1^. It predominantly affects children and young adults, with peak incidence between ages 10 and 25 ^2, 3^. The standard treatment—neoadjuvant chemotherapy, surgical resection, and adjuvant chemotherapy—cures approximately two-thirds of patients with localized disease ^4^. However, survival rates have plateaued since the 1980s ^5^, largely due to intrinsic chemoresistance and remarkable inter- and intratumoral heterogeneity ^6^. This complexity, particularly within the tumor microenvironment, remains poorly understood and represents a critical barrier to improving clinical outcomes.

Recent advances in single-cell RNA sequencing (scRNA-seq) have provided high-resolution insights into the diverse cellular ecosystem of OS, surpassing the capabilities of bulk transcriptomic profiling ^7^. Previous scRNA-seq studies have highlighted cellular heterogeneity, immune suppression, and tumor–stromal interactions in OS, albeit with limited sample sizes and a lack of spatial context. For instance, Zhou et al. profiled 11 post-chemotherapy samples, revealing intratumoral heterogeneity, immunosuppressive landscapes, and malignant cell-stromal interactions ^8^. Liu et al. provided a baseline profile using 6 treatment-naïve samples ^9^. While these studies, including our own initial atlas ^10^, advance understanding of disease mechanisms and potential therapeutic targets, significant limitations persist. The collective dataset remains small compared to other malignancies, restricting comprehensive TME characterization. More critically, the spatial architecture of the OS tumor microenvironment (TME)—including the distribution, co-localization, and interaction of malignant and non-malignant cells—remains largely unexplored. Understanding this spatial organization is essential for decoding functional heterogeneity and identifying niche-specific therapeutic vulnerabilities.

Here, we present a comprehensive integrated analysis of one of the largest osteosarcoma cohorts to date, leveraging paired scRNA-seq and spatial transcriptomics complemented by multiplexed immunofluorescence (mIF) validation. We constructed a cellular meta-map to decode TME heterogeneity across patients with diverse ages, stages, and treatment histories. Our study reveals distinct, spatially resolved TME niches: Chondroblastic osteosarcoma localizes to vascular-depleted, hypoxic regions, driven by hypoxia signaling pathways and exhibiting reduced immune infiltration; Osteoblastic osteosarcoma localizes within highly vascularized niches characterized by an enhanced immunosuppressive microenvironment. Within the osteoblastic niche, we identify key mechanistic pathways: Osteoblastic tumor cells activate cancer-associated fibroblasts (CAFs) via SPP1 signaling; Activated CAFs appear to modulate T cell activity through myeloid intermediates; Osteoblastic regions are enriched in osteoclasts, potentially regulated by tumor-derived BMP signaling to promote maturation. This work defines the spatial and cellular architecture of the human osteosarcoma TME, uncovering novel niche-specific mechanisms driving heterogeneity. Our findings provide a framework for understanding TME-driven therapy resistance and reveal promising targets to broaden therapeutic efficacy.

## Results

### A High-resolution Cellular Landscape of Human Osteosarcomas

The overall workflow of this study was illustrated in Fig 1A. We generated single-cell transcriptomes from 14 human osteosarcoma samples (Table 1), including 8 post-chemotherapy specimens (7 primary tumors and 1 recurrent tumor) and 6 treatment-naïve primary tumors. After rigorous quality control filtering (see Methods), a total of 78,838 high-quality single cells (median genes detected per cell: 1,795) were retained for downstream analysis using the Seurat pipeline. Uniform Manifold Approximation and Projection (UMAP) visualization revealed distinct cellular clusters primarily driven by lineage identity (Figure 1B). Cells expressing canonical lineage markers—including myeloid cells (LYZ, CD68), T and natural killer (NK) lymphocytes (CD3D, NKG7), and endothelial cells (VWF, PECAM1)—consistently clustered across different patients (Figures 1B and 1C). In contrast, cells of mesenchymal origin exhibited two distinct clustering patterns: (1) cross-patient clusters primarily consisting of fibroblasts and pericytes, and (2) patient-specific clusters. InferCNV analysis confirmed that cross-patient mesenchymal clusters corresponded to normal stromal populations, such as fibroblasts and pericytes (Figure S2I). Notably, the close transcriptional similarity between osteosarcoma cells and fibroblasts complicated their separation using batch-correction tools such as Harmony (Figure S1A), underscoring a potential source of misclassification in previous single-cell studies.

**Figure 1.**
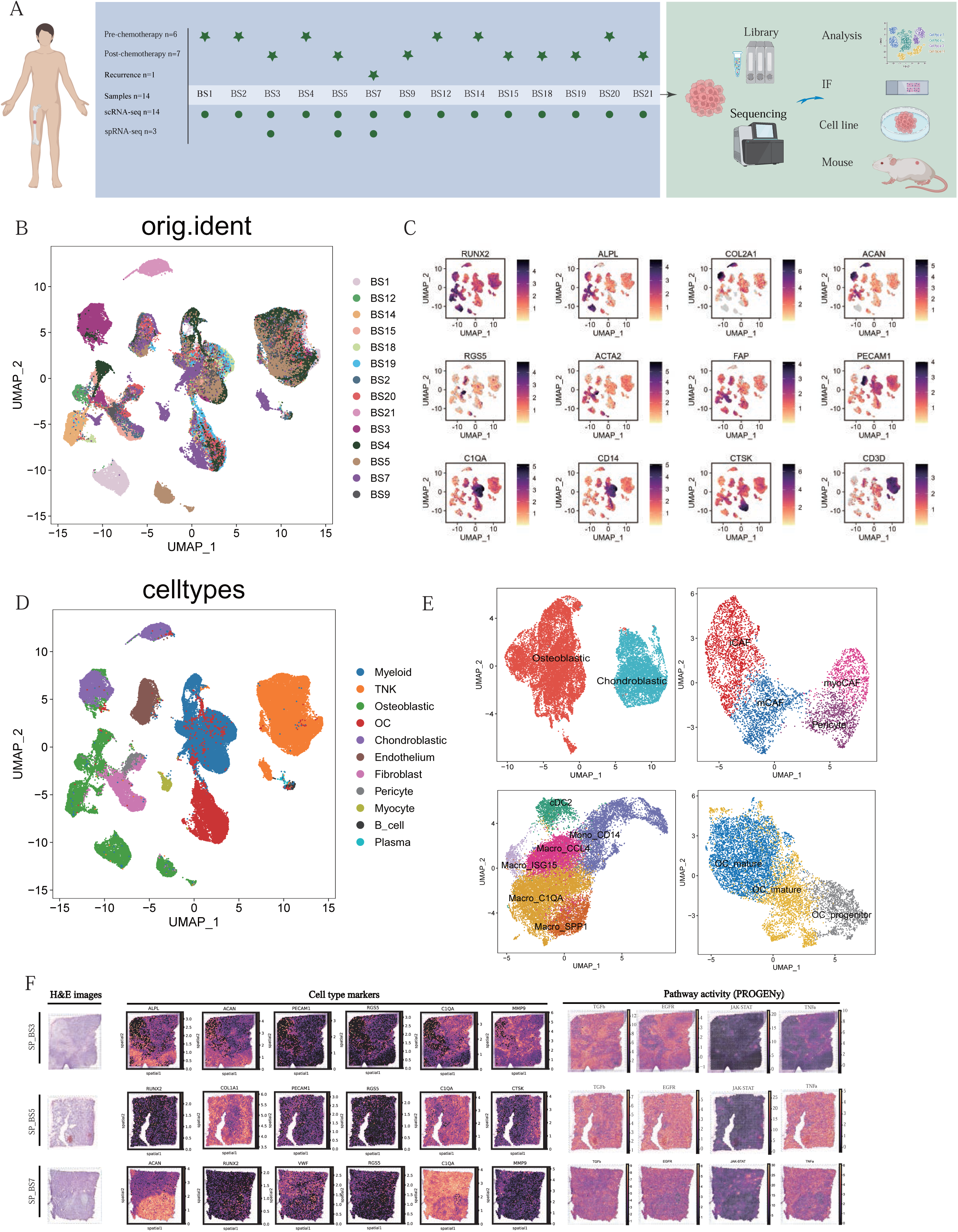
Single-cell and spatial atlas of the osteosarcoma tumor microenvironment. (A) Schematic overview of the study design. (B) UMAP embedding of 78,838 high-quality single cells from 14 osteosarcoma samples. (C) Expression of canonical lineage markers (e.g., CD3D, LYZ, PECAM1) confirming robust clustering of immune and endothelial populations across patients. (D) Identification of 11 major cellular subtypes within the tumor microenvironment (TME), including osteoblastic and chondroblastic tumor cells, immune cells (myeloid, T/NK, B, plasma), fibroblasts, endothelial cells, osteoclasts, and myocytes. (E) Subclustering of mesenchymal, myeloid, and osteoclast lineages. Tumor cells segregate into osteoblastic (RUNX2, ALPL) and chondroblastic (SOX9, ACAN, COL2A1) lineages. Fibroblasts are further subclassified into myoCAFs, mCAFs, iCAFs, and pericytes; myeloid cells into six subsets; osteoclasts into three maturation states. (F) Representative spatial transcriptomic slides overlaid with H&E staining, expression of canonical cell-type marker genes, and inferred pathway activity profiles.

**Figure S1.**
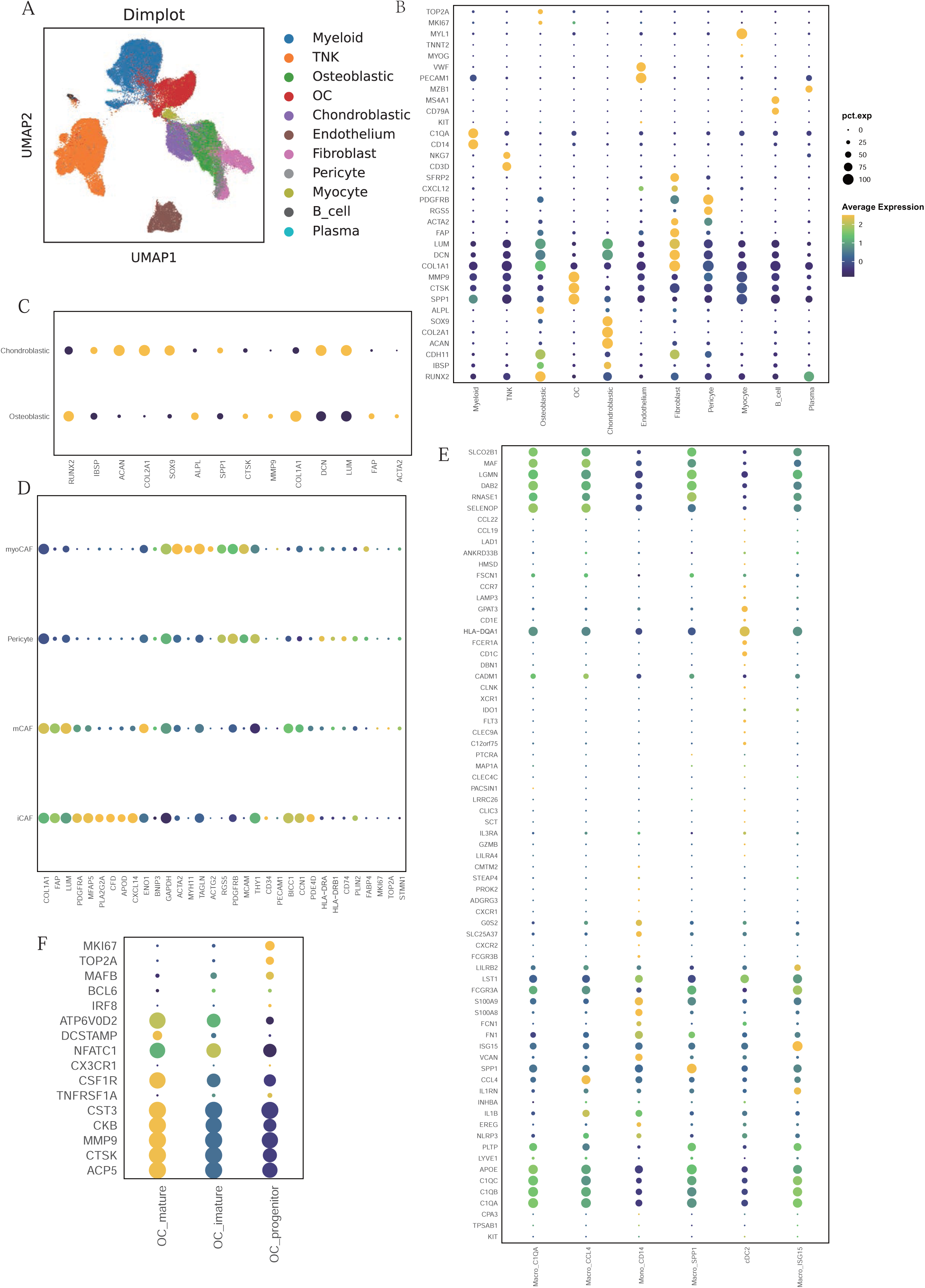
Cell type marker expression and batch correction validation. **(A)** UMAP plot after batch correction, demonstrating correction of patient-specific effects while preserving cell-type identity. **(B)** Dotplot showing the expression of canonical marker genes used for major cell type annotation. **(C)** Dotplot confirming osteoblastic (ALPL, RUNX2) and chondroblastic (SOX9, ACAN) marker expression in tumor clusters. **(D)** Dotplot for fibroblast subtypes: myoCAFs (ACTA2, TAGLN), matrix CAFs (COL1A1, FAP), iCAFs (CXCL14, CFD), and pericytes (RGS5, PDGFRB). **(E)** Dotplot for six myeloid subclusters, including SPP1+ macrophages and ISG15+ interferon-stimulated macrophages. **(F)** Dotplot for osteoclast subclusters.

Based on established marker gene expression, we annotated 11 major cellular subtypes within the tumor microenvironment (Figure 1D, Fig S1B, Table 2): osteoblastic osteosarcoma cells (Osteoblastic, n=13,404) characterized by RUNX2 and ALPL expression; chondroblastic osteosarcoma cells (Chondroblastic, n=6,091) expressing ACAN, SOX9, and COL2A1 ^8^; myeloid cells (Myeloid, n=19,868) marked by CD14 and C1QA]; T and NK lymphocytes (TNK, n=19,802) defined by CD3E and NKG7 expression ^11^; fibroblasts (Fibroblast, n=4,527) positive for COL1A1 and ACTA2 ^12^; endothelial cells (Endothelium, n=5,554) expressing PECAM1 and VWF ^12^; osteoclasts (OC, n=8,395) characterized by CTSK and MMP9 ^13^; B cells (n=466) marked by MS4A1 and CD79A ^12^; plasma cells (Plasma, n=132) expressing MZB1 ^14^; and myocytes (Myocyte, n=740) identified by MYOG ^15^.

Further subset analyses of tumor cells revealed that osteosarcoma cells segregate primarily into two distinct lineages reflecting clinical phenotypes: an osteoblastic lineage (RUNX2, ALPL) and a chondrocytic lineage (ACAN, SOX9, COL2A1) (Fig 1E, Fig S1C). Normal mesenchymal cells were resolved into four clusters: myofibroblastic cancer-associated fibroblasts (myoCAFs; ACTA2, TAGLN), pericytes (RGS5, PDGFRB), matrix cancer-associated fibroblasts (mCAFs; COL1A1, FAP), and inflammatory cancer-associated fibroblasts (iCAFs; CFD, CXCL14) (Fig 1E, Fig S1D). Within the myeloid compartment, six distinct clusters were identified: Macro_C1QA (C1QA, APOE), Macro_CCL4 (CCL4, IL1B), Mono_CD14 (S100A8, S100A9), Macro_SPP1 (SPP1), cDC2 dendritic cells (HLA-DQA1), and Macro_ISG15 (ISG15) (Fig 1E, Fig S1E). Osteoclast populations were stratified into three maturation states: progenitors (OC_progenitor; MKI67, TOP2A), immature osteoclasts (OC_immature; moderate expression of ACP5, CTSK, MMP9, CKB, CST3), and mature osteoclasts (OC_mature; highest expression levels of ACP5, CTSK, MMP9, CKB, CST3) (Fig 1E, Fig S1F).

To investigate spatial heterogeneity, we performed spatial transcriptomics (ST) analysis on samples BS3, BS5, and BS7. Samples SP_BS3 and SP_BS5 were derived from post-chemotherapy osteosarcoma tissues, while SP_BS7 represented a post-chemotherapy recurrent tumor. Consistent with the known histopathological features of osteosarcoma, hematoxylin and eosin (H&E) staining and transcriptomic profiling revealed no clear-cut boundaries between tumor and stromal compartments (Fig 1F). Instead, spatial analysis identified osteogenic status as the predominant organizer of the tumor microenvironment. Regions were broadly categorized as ossified (mineralized) or poorly ossified, suggesting that osteogenesis may be a primary determinant of TME architecture. We mapped the spatial distribution of key cell subtypes identified by scRNA-seq using their canonical marker genes. Pathway enrichment analysis via PROGENy revealed distinct spatial signaling patterns: TGF-βsignaling was enriched predominantly within osteoblastic, chondroblastic, and fibroblast-rich regions; EGFR signaling localized to vascular-enriched areas; JAK-STATpathway activity was prominent in osteoclast-rich regions, implicating its role in osteoclast function; TNFαsignaling was enriched in myeloid cell-dense zones.

In summary, this study provides a comprehensive single-cell and spatial transcriptomic atlas of human osteosarcoma, delineating the major cellular constituents and their spatial organization within the TME. Our integrated analysis, supported by histological validation, identifies osteogenic status as a key driver of spatial heterogeneity and lays the groundwork for future mechanistic investigations and potential therapeutic targeting.

### Hypoxia-Driven Metabolic Reprogramming Promotes Chondroblastic Trans-Differentiation in Osteosarcoma

To investigate lineage-specific heterogeneity in osteosarcoma, we projected annotated cell types onto the pre-batch-correction UMAP embedding. Strikingly, within individual patients, osteoblastic and chondroblastic tumor cells clustered together rather than segregating by differentiation state (Fig 2A, Fig S2A)), suggesting that inter-patient heterogeneity outweighs intra-tumoral variation. Chondroblastic tumor cells exhibited markedly reduced mitochondrial gene expression compared to their osteoblastic counterparts (Fig. S2B), indicative of suppressed metabolic activity. This phenotype was corroborated by decreased proliferative potential, as evidenced by lower MKI67 expression and diminished representation in proliferative cell cycle phases (Fig. S2C-D). These findings collectively define a metabolically quiescent state in chondroblastic osteosarcoma cells. We further observed that osteoblastic cells predominated in treatment-naïve samples, whereas chondroblastic cells were more abundant in post-chemotherapy samples (Fig. S2E), implying that chemotherapeutic stress may drive chondroblastic trans-differentiation.

**Figure 2.**
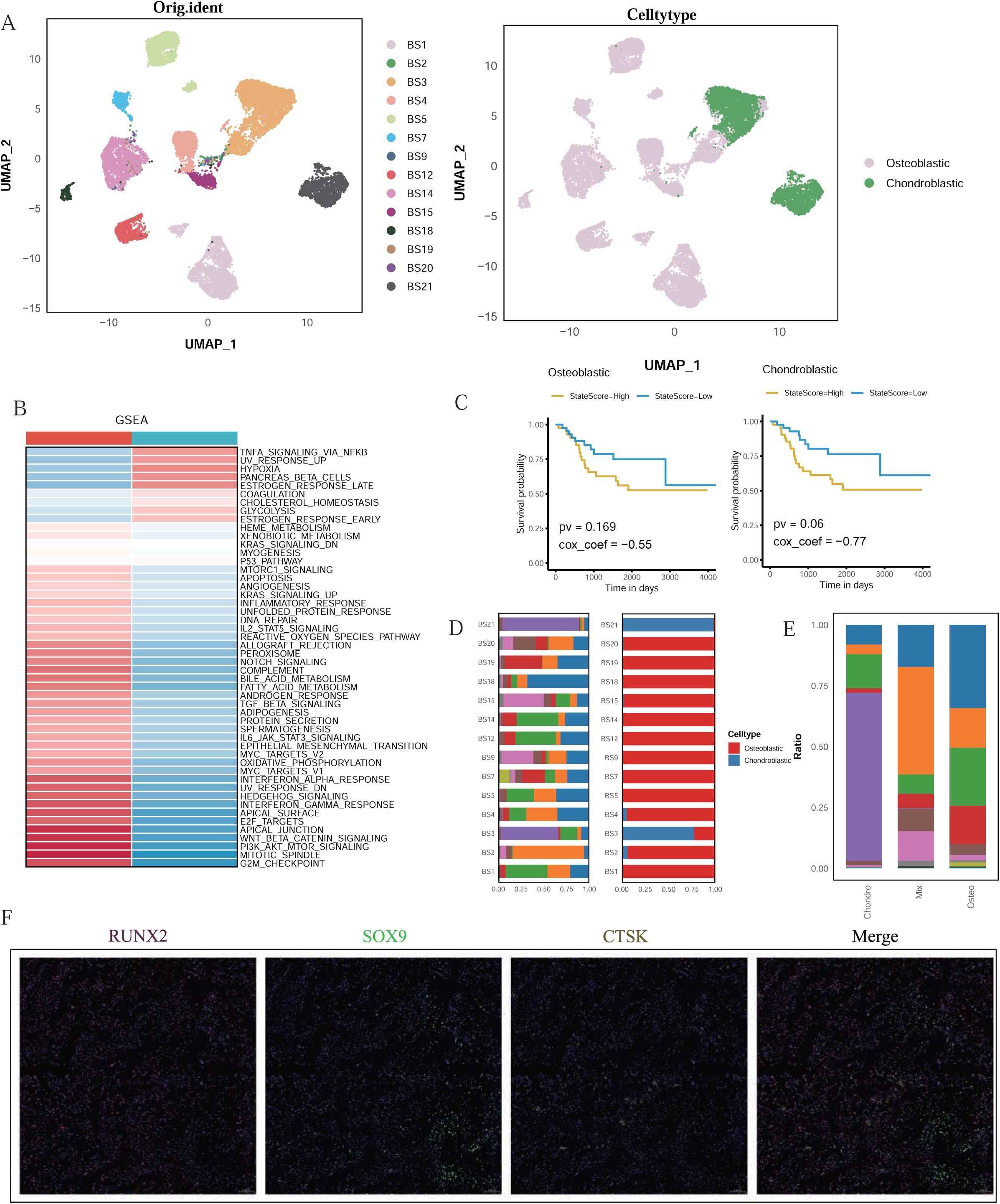
Hypoxia drives trans-differentiation of osteosarcoma cells toward a chondroblastic state. (A) UMAP visualization of annotated tumor cells overlaid on the pre-batch-correction embedding. Osteoblastic and chondroblastic tumor cells clustered more by patient than by lineage, indicating stronger inter-tumor than intra-tumor heterogeneity. (B) GSVA pathway enrichment scores for osteoblastic and chondroblastic tumor cells. (C) Kaplan–Meier survival curves showing poorer prognosis in patients with tumors enriched for osteoblastic or chondroblastic gene signatures. (D–E) Barplots comparing TME composition in osteoblastic vs. chondroblastic niches. Osteoblastic tumors harbored more CAFs, vasculature, T/NK cells, and osteoclasts, while chondroblastic regions showed sparse stromal and immune infiltration. (F) Multiplex immunofluorescence staining of sample BS3 revealed preferential localization of osteoclasts to osteoblastic regions.

**Figure S2.**
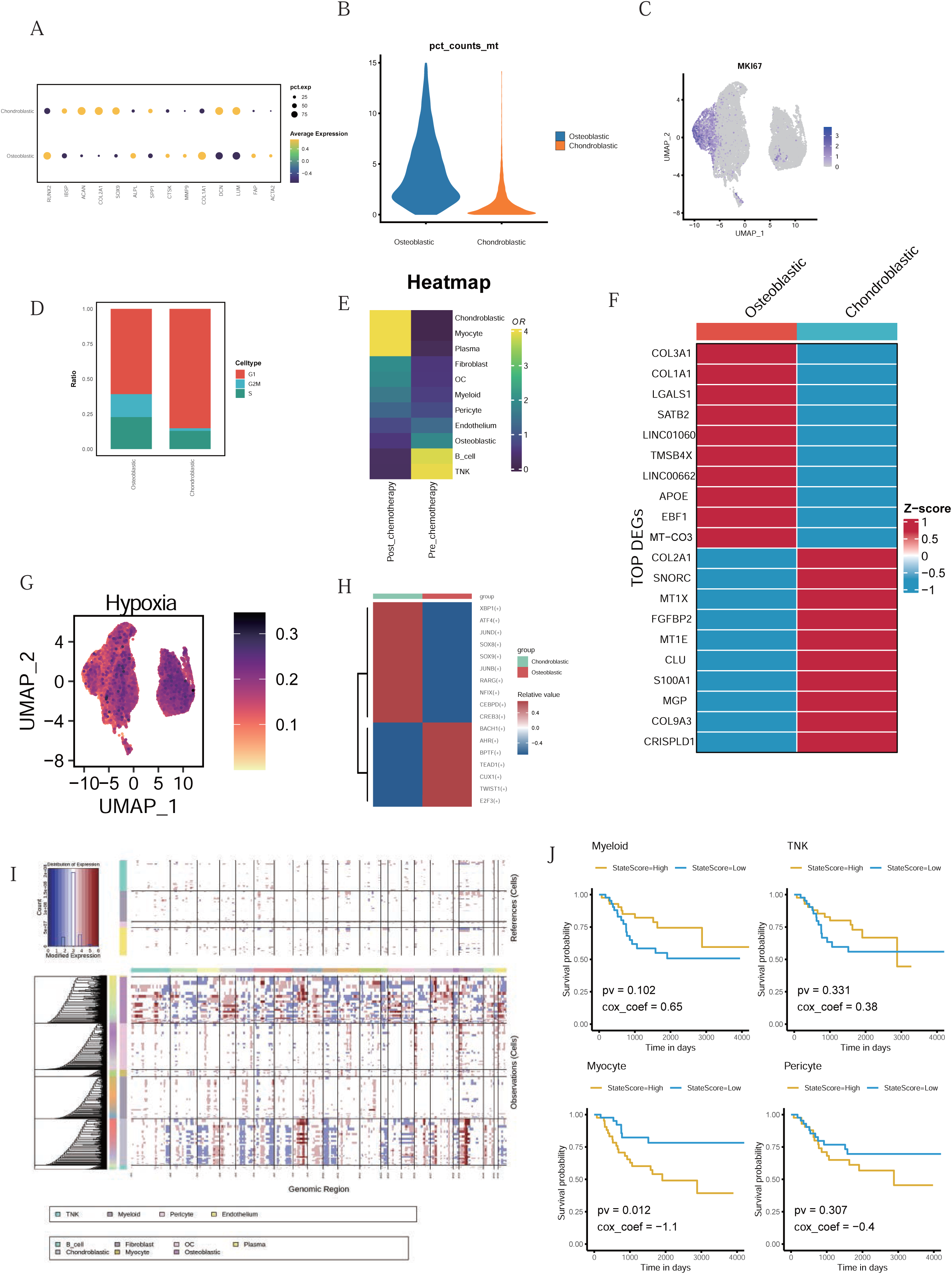
Molecular and functional features of osteoblastic and chondroblastic tumor lineages. **(A)** Dot plot showing the expression of canonical marker genes distinguishing osteoblastic and chondroblastic osteosarcoma cell populations. **(B)** Violin plot showing elevated mitochondrial gene content in osteoblastic cells. **(C–D)** MKI67 expression and inferred cell cycle states, highlighting higher proliferative activity in osteoblastic cells. **(E)** Heatmap showing the relative abundance of osteoblastic and chondroblastic tumor cells across treatment-naïve and post-chemotherapy osteosarcoma samples. **(F)** Heatmap of top differentially expressed genes between osteoblastic and chondroblastic cells. **(G)** Featureplot showing hypoxia pathway activity scores in osteoblastic and chondroblastic tumor cells. **(H)** SCENIC-inferred transcription factor activity showing enrichment of SOX family TFs (e.g., SOX9) in chondroblastic cells. **(I)** CNV heatmap revealing distinct genomic alterations between the two tumor lineages. **(J)** Kaplan–Meier survival curves showing better prognosis in patients with tumors enriched for T/NK or myeloid gene signatures.

Differential gene expression analysis revealed that osteoblastic cells were enriched for matrix-associated genes such as COL1A1, COL3A1, and APOE, whereas chondroblastic cells preferentially expressed MGP, CLU, and COL9A3 (Fig. S2F, Table3). Notably, MGP functions to inhibit pathological calcification by maintaining extracellular calcium solubility, implicating chondroblastic differentiation in disrupted osteogenesis ^16^. Gene set variation analysis (GSVA) revealed that chondroblastic tumor cells display hallmarks of metabolic suppression, characterized by attenuated oxidative phosphorylation and enhanced glycolytic dependency, whereas osteoblastic cells showed enrichment in epithelial-mesenchymal transition (EMT), angiogenesis, and interferon-gamma response pathways (Fig 2B, Fig S2G, Table 4). SCENIC-based transcription factor activity analysis highlighted upregulation of the SOX family in chondroblastic cells, supporting a model wherein HIF1A-mediated hypoxic signaling induces SOX9, thereby promoting chondrogenic reprogramming (Fig S2H). This aligns with our previous findings confirming co-localization of HIF1A and SOX9 in osteosarcoma ^17^. Further, copy number variation (CNV) profiling revealed distinct chromosomal alterations between osteoblastic and chondroblastic tumor populations (Fig. S2I), underscoring their molecular divergence beyond transcriptional and metabolic reprogramming.

Given the clinical heterogeneity of osteosarcoma, we hypothesized that differences in tumor microenvironment composition may underlie variable prognoses. Survival analysis based on gene expression signatures of osteoblastic and chondroblastic cells revealed that both differentiation states were associated with poorer overall survival, whereas T/NK lymphocyte and myeloid cell signatures correlated with improved prognosis (Fig 2C, Fig S2J, Table 5-7). These data indicate that tumor purity and immune infiltration impact clinical outcomes, with chondroblastic differentiation linked to particularly unfavorable prognosis.

Consistent with these findings, osteoblastic osteosarcomas exhibited higher immune infiltration, including T/NK cells, myeloid cells, and osteoclasts. Conversely, chondroblastic osteosarcomas, enriched in hypoxia-related pathways, displayed reduced vascular components such as endothelial cells and pericytes within the TME (Fig 2D, E, Table 8-9). To validate our single-cell observations, we performed immunofluorescence staining for osteoclast markers in sample BS3, which contains both osteoblastic and chondroblastic regions. Osteoclasts preferentially localized to osteoblastic areas (Figure 2F), supporting our computational inferences.

Collectively, these data position hypoxia-induced chondroblastic trans-differentiation as a convergence point for metabolic dysregulation, therapy resistance, and immune evasion—a triad that may be therapeutically targeted via HIF1A inhibition or vascular normalization strategies.

### Spatially Resolved Cellular Niches Underlie Osteosarcoma Heterogeneity

To investigate the spatial architecture of osteosarcoma tissues, we leveraged spatial transcriptomic profiling. Given that each spatial spot captures transcriptomic signals from multiple neighboring cells, we performed spot-level deconvolution using matched single-cell RNA-seq data to infer the cellular composition within each spot (Fig. 3A). The deconvolution results showed high concordance with the single-cell annotations, supporting the validity of our approach.

**Figure 3.**
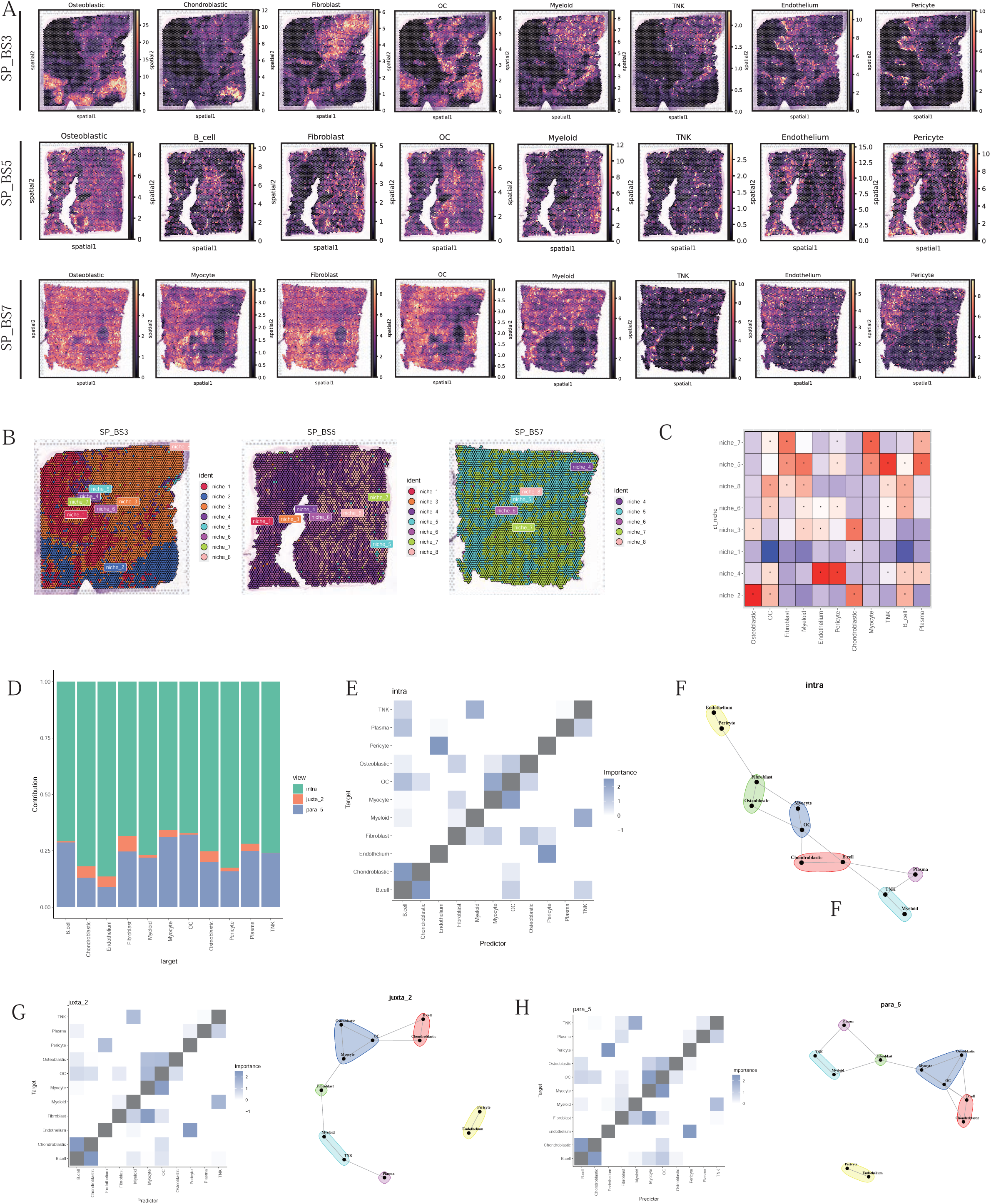
Spatially defined niches and coordinated cell-type interactions in osteosarcoma. **(A)** Estimated cell-type composition across spatial transcriptomic spots based on Cell2location deconvolution using matched scRNA-seq reference data. **(B)** Unsupervised clustering of spatial spots revealed eight conserved niches across three ST slides. **(C)** Heatmap of cell-type enrichments within each niche. **(D)** MISTy modeling showed that intra-spot colocalization was the strongest predictor of local cell-type abundance. **(E)** Median intraview importance scores showing the predictive influence of each cell type on the abundance of other cell types within the same Visium spot, as inferred by MISTy. **(F)** Schematic representation of intraview cell–cell interaction networks within Visium spots. **(G–H)** Median juxtaview (adjacent spot) and paraview (extended neighborhood) importance scores representing the predictive impact of spatially neighboring cell types on local cell-type abundance.

**Figure S3.**
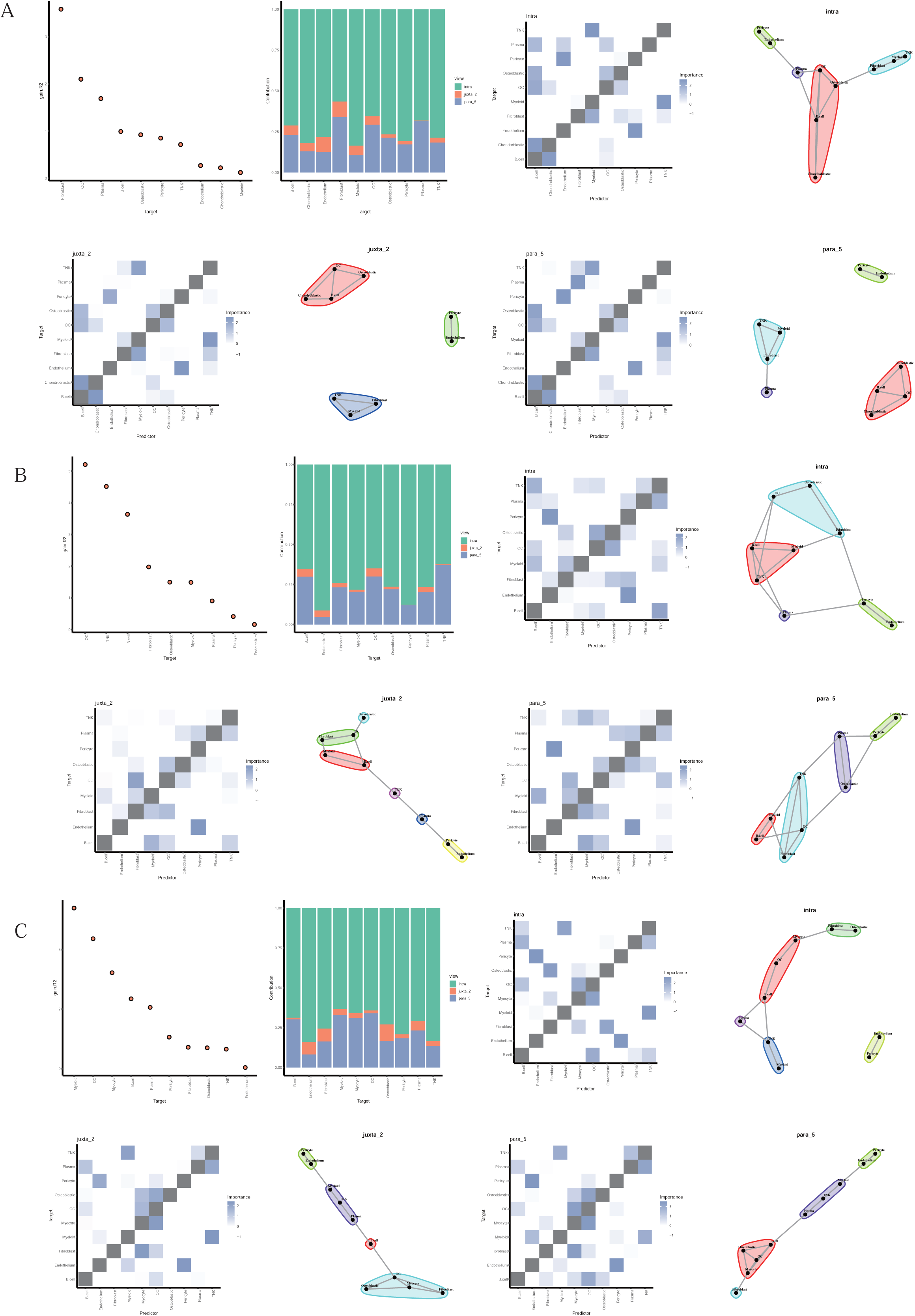
Sample-specific spatial co-localization patterns. **(A–C)** Spatial co-localization analysis of osteosarcoma samples SP_BS3, SP_BS5, and SP_BS7 performed separately. Cell–cell interaction patterns were modeled across intraview (within-spot), juxtaview (adjacent-spot), and paraview (extended neighborhood) spatial contexts, revealing consistent and spatially dependent associations among cell types across tumor regions.

Unsupervised clustering based on cell-type proportions across all spatial spots identified eight distinct spatial niches, which likely represent conserved structural and functional units across tumors (Fig. 3B). Spatial mapping of these niches revealed substantial inter-tumoral variation: for example, SP_BS3 was predominantly composed of niches 1, 2, and 3; SP_BS5 was enriched in niches 4, 6, and 8; while SP_BS7 was largely occupied by niches 5 and 7. This observation highlights the considerable spatial heterogeneity across osteosarcoma specimens. We next mapped annotated cell types from the single-cell data to these spatial niches to infer their cellular identities (Fig. 3C). In SP_BS7, two anatomically distinct tumor regions—the ossified and incompletely ossified areas—corresponded to niche 5 and niche 7, respectively. Niche 5 was enriched in myeloid and T/NK cells and localized to the ossified region, whereas niche 7 was characterized by fibroblast and myocyte enrichment, aligning with regions of incomplete ossification. Notably, niche 2 consisted primarily of osteoblastic, chondroblastic, and osteoclast populations, while niche 4 was enriched in vascular components, including endothelial cells and pericytes. These findings underscore that osteosarcoma tissues are composed of spatially segregated cellular ecosystems with distinct functional specializations. Spatial co-localization analysis revealed that osteoblastic and chondroblastic cells frequently co-occurred within niches 2 and 3, rather than forming discrete spatial compartments. This pattern suggests that these differentiation states are not derived from independent tumor clones, but instead represent phenotypic plasticity shaped by shared microenvironmental cues, particularly hypoxia.

To quantify how spatial context influences local cell-type composition, we employed the MISTy framework to model predictive relationships at three spatial resolutions: (1) within individual spots (colocalization), (2) within adjacent spots (local neighborhood, radius = 1), and (3) within extended neighborhoods (radius = 15 spots). Across tumor regions, intraview (within-spot) colocalization provided the strongest predictive signal for cell-type abundance (Fig. 3D). Endothelial cells were consistently the most predictive determinant of pericyte abundance across all spatial scales (intra-, juxta-, and paraview), reflecting their known perivascular association (Fig. 3E–F). Similarly, T/NK cells and myeloid populations exhibited strong mutual dependencies, corresponding to immune-rich niches (e.g., niches 5, 6, and 8). Additionally, myeloid cells and fibroblasts showed robust co-enrichment, particularly within niches 5 and 8, consistent with the role of fibroblasts in orchestrating macrophage recruitment and activation. These spatial dependencies remained stable across both immediate and extended neighborhoods (Fig. 3G–H), and exhibited generally consistent patterns across individual samples (Fig. S3A–C)

To functionally link spatial organization with molecular signaling programs, we assessed the spatial association between key signaling pathways and cell types. As anticipated, chondroblastic cells were spatially enriched in regions exhibiting high hypoxia pathway activity (Fig. S4A). Although chondroblastic cells were not annotated in the single-cell dataset for SP_BS7 (likely due to sampling bias), spatial transcriptomic analysis revealed that areas of incomplete ossification in this sample lacked vascularization and exhibited elevated expression of hypoxia-associated genes, including the chondroblastic marker ACAN (Fig. S4B).

In summary, despite extensive inter- and intra-tumoral cellular heterogeneity, osteosarcoma tissues exhibit reproducible spatial co-localization patterns. These spatially organized microenvironments appear to govern differentiation trajectories and immune infiltration. In particular, hypoxia emerges as a central driver of chondroblastic transdifferentiation and spatial immune exclusion, suggesting that microenvironmental modulation—such as targeting hypoxia or normalizing vasculature—may hold therapeutic potential in osteosarcoma.

### Spatially Resolved Molecular Niches and Fibroblast-Centric Signaling in Osteosarcoma

To delineate the molecular heterogeneity of osteosarcoma in a spatially resolved and unbiased fashion, we performed unsupervised clustering of spatial transcriptomic spots based on gene expression profiles, which revealed a set of distinct molecular niches (Fig. 4A–B). Spatial mapping of these niches demonstrated strong concordance with histological structures observed via H&E staining, particularly in sample SP_BS7. For instance, niche 0 co-localized with the osteogenic region, while niche 2 corresponded to areas of incomplete ossification. Niche 12 was defined by high expression of COL2A1 and COL11A1, suggestive of a chondrogenic identity, whereas niche 11 was enriched for HBA1/HBA2 and HBB, indicative of a hematopoietic compartment (Fig. S4C). These data suggest that molecular niches represent spatially distinct and functionally specialized microenvironments within osteosarcoma.

**Figure 4.**
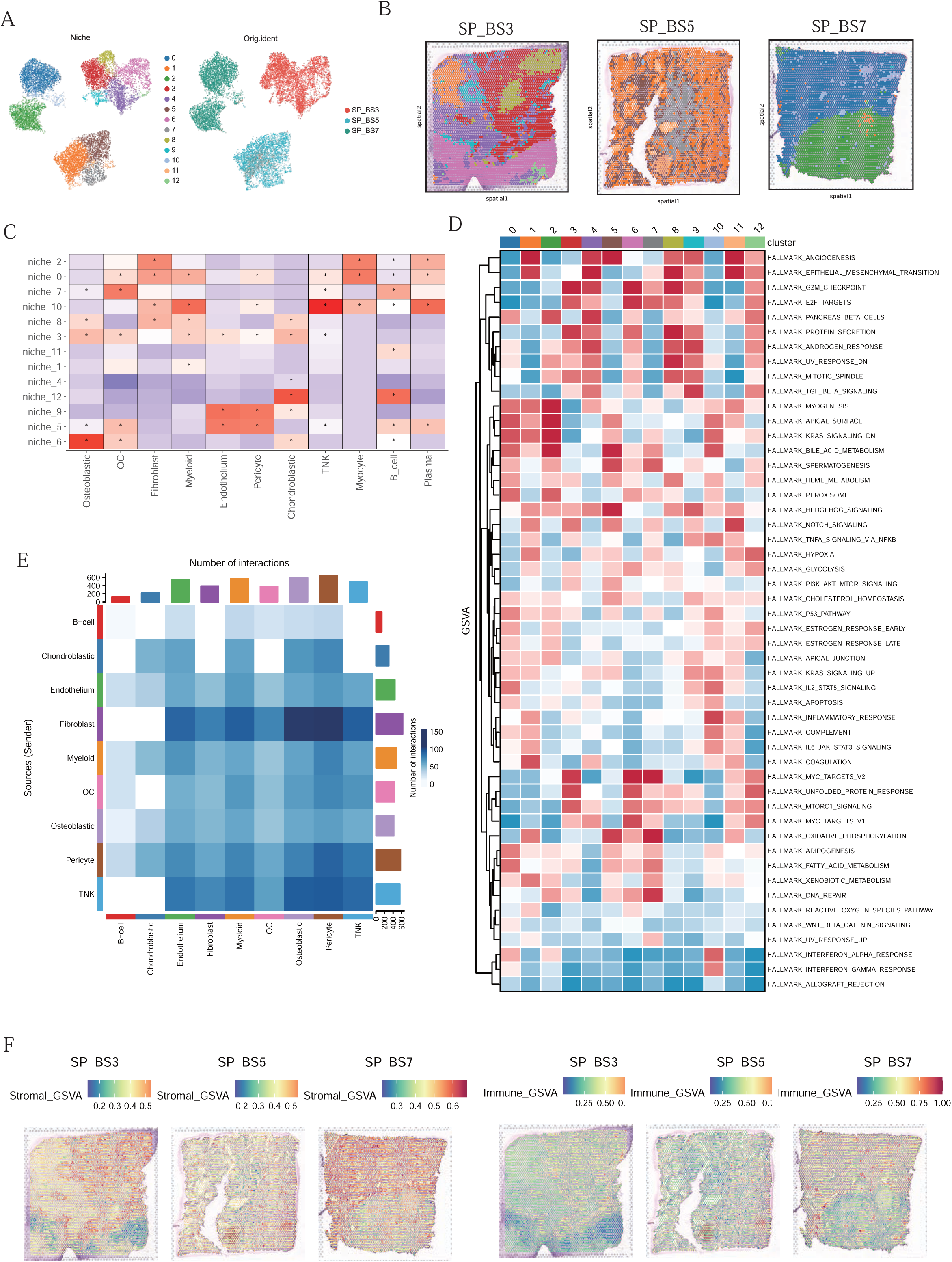
Molecularly distinct spatial niches reflect underlying cellular and functional diversity. **(A–B)** Unsupervised clustering of spatial transcriptomic spots based on gene expression identified 13 molecular niches. **(C)** Heatmap of cell-type abundance across niches, highlighting niche-specific enrichment of osteoblastic, chondroblastic, immune, stromal, and vascular components. **(D)** GSVA analysis of hallmark pathways reveals niche-specific signaling patterns. **(E)** CellChat-inferred signaling networks show fibroblasts as central hubs interacting with osteoblastic, immune, and myeloid cells. **(F)** Co-localization of stromal and immune pathway scores supports a role for fibroblasts in immune landscape modulation.

**Figure S4.**
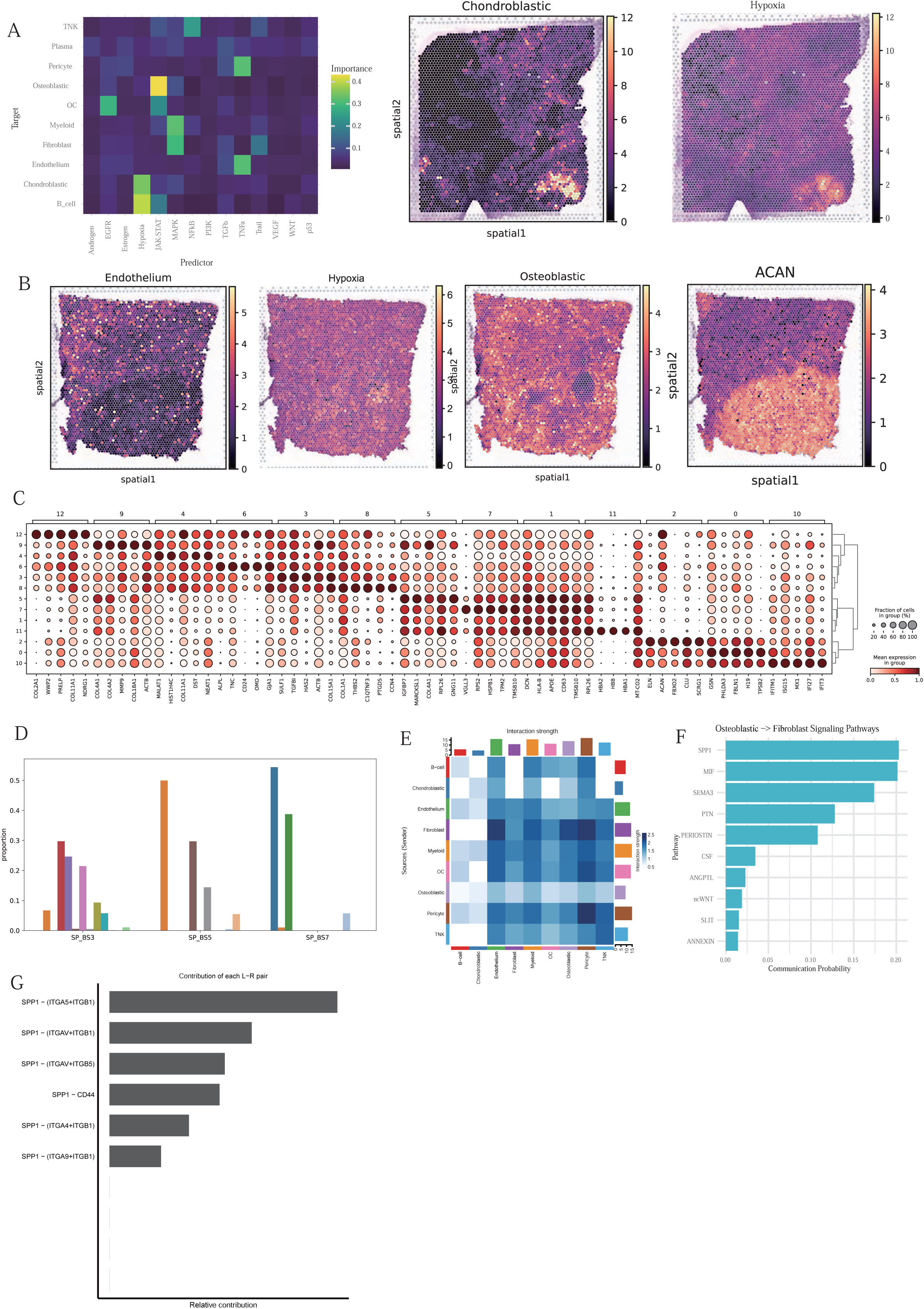
Spatial co-localization of signaling pathways and niche identity. **(A)** Spatial distribution of hypoxia pathway activity in sample SP_BS3, showing co-localization with regions enriched in chondroblastic tumor cells. **(B)** Spatial expression of chondrocytic marker ACAN in hypoxic, poorly vascularized zones of SP_BS7. **(C)** Expression of niche-specific marker genes (e.g., COL2A1, HBB) supporting molecular niche identity. **(D)** Proportions of molecular niches across the three ST slides. **(E)** Interaction strength between fibroblasts and immune cell populations, inferred from CellChat analysis. **(F)** SPP1-mediated signaling is predicted to be the most active pathway from osteoblastic tumor cells to fibroblasts. **(G)** Contribution of individual ligand–receptor pairs within the SPP1 signaling pathway from osteoblastic tumor cells to fibroblasts.

Integrative analysis across three osteosarcoma specimens identified 13 molecular niches, underscoring pronounced inter-tumoral diversity. SP_BS3 was dominated by niches 3, 4, and 6; SP_BS5 was enriched for niches 1, 5, and 7; and SP_BS7 primarily consisted of niches 0 and 2 (Fig. S4D). Notably, niche 1 was the only molecular niche shared across all three samples, emphasizing both the extensive molecular diversity and the presence of a potentially conserved tumor microenvironmental element in osteosarcoma. To investigate cellular underpinnings of molecular diversity, we analyzed spot-level cell-type composition using Cell2location-based deconvolution. Cell-type abundances varied markedly across niches both within and across tumors. Co-analysis of molecular states and cellular compositions revealed that niche 0 was enriched in osteoclasts, fibroblasts, myeloid cells, T/NK cells, and myocytes, reflecting a complex immune–stromal microenvironment. In contrast, niches such as 1, 4, and 11 were composed of fewer cell types but exhibited highly distinct transcriptional signatures. Niche-specific cellular enrichment patterns included endothelial–pericyte dominance in niches 5 and 9, osteoblast–osteoclast co-localization in niche 6, and chondroblastic predominance in niche 12 (Fig. 4C). These findings demonstrate that molecular niches are closely shaped by underlying cellular architecture.

To functionally annotate the spatial niches, we performed Gene Set Variation Analysis using the Hallmark gene sets (Fig. 4D, Table 10). Distinct functional profiles emerged across niches, reflecting regional specialization. Niche 12, dominated by chondroblastic cells, showed robust enrichment for hypoxia-related pathways. Niche 10 exhibited signatures of inflammatory response and interferon signaling, while niche 1 was strongly associated with angiogenesis. Additional niches displayed pathway-specific enrichments, such as myogenesis and epithelial signatures in niche 2, Hedgehog signaling in niche 5, oxidative phosphorylation in niche 7, and TGF-β signaling in niche 9. These spatially distinct pathway landscapes further underscore the transcriptional and functional heterogeneity of osteosarcoma.

To dissect intercellular communication across spatial niches, we applied CellChat to infer ligand–receptor signaling networks. Fibroblasts emerged as central signaling hubs, exhibiting extensive interactions with osteoblastic, myeloid, and T/NK cells (Fig. 4E, S4E). Supporting this observation, GSVA revealed strong spatial co-localization between stromal and immune scores (Fig. 4F), reinforcing the concept that cancer-associated fibroblasts (CAFs) actively shape the immunological landscape of the osteosarcoma microenvironment. Further analysis identified SPP1 as a key ligand mediating interactions between osteoblasts and fibroblasts (Fig. S4F), implicating this pathway in CAF activation and broader stromal remodeling. We next delineated a fibroblast-centered communication axis influencing immune regulation and bone remodeling. Fibroblasts were predicted to modulate macrophage populations via MIF and ANXA1 signaling, while myeloid cells influenced T/NK cells through LGALS9 (Fig. S5A). Although osteoclasts originate from the myeloid lineage, spatial co-localization analysis revealed that their distribution was distinct from that of myeloid cells across individual samples. Instead, osteoclasts consistently localized in proximity to osteoblastic cells, suggesting that osteoblast-derived signals may play a more direct role in guiding osteoclast recruitment and spatial positioning. Cell–cell communication modeling further identified osteoblasts as potential regulators of osteoclast activity through suppression of MIF and activation of BMP and SEMA family ligands (Fig. S5B). Supporting this interaction axis, pseudotime trajectory analysis revealed progressive upregulation of osteoclast maturation markers—including ACP5, CKB, CTSK, and MMP9—consistent with osteoblast-mediated promotion of osteoclast differentiation (Fig. S5C–D). Collectively, these findings reveal that spatially defined molecular niches in osteosarcoma are shaped by distinct cellular compositions and coordinated intercellular communication networks, with fibroblasts serving as key modulators of immune and bone-related processes.

### SPP1 Promotes Osteosarcoma Cell Proliferation and Activates Cancer-Associated Fibroblasts

To investigate the role of SPP1 in osteosarcoma progression, we first examined its expression across multiple tumor types (http://timer.cistrome.org/) and found that SPP1 is broadly overexpressed in various cancers (Fig S6A). In 143B osteosarcoma cells treated with low-dose doxorubicin, bulk RNA sequencing (n = 3 pairs) revealed a significant upregulation of SPP1 following treatment (Fig. 5A). Gene Ontology (GO) enrichment analysis of differentially expressed genes indicated activation of pathways involved in extracellular matrix organization, extracellular structure organization, and cellular respiration, implicating SPP1 in both stromal remodeling and metabolic regulation (Fig. 5B).

**Figure 5.**
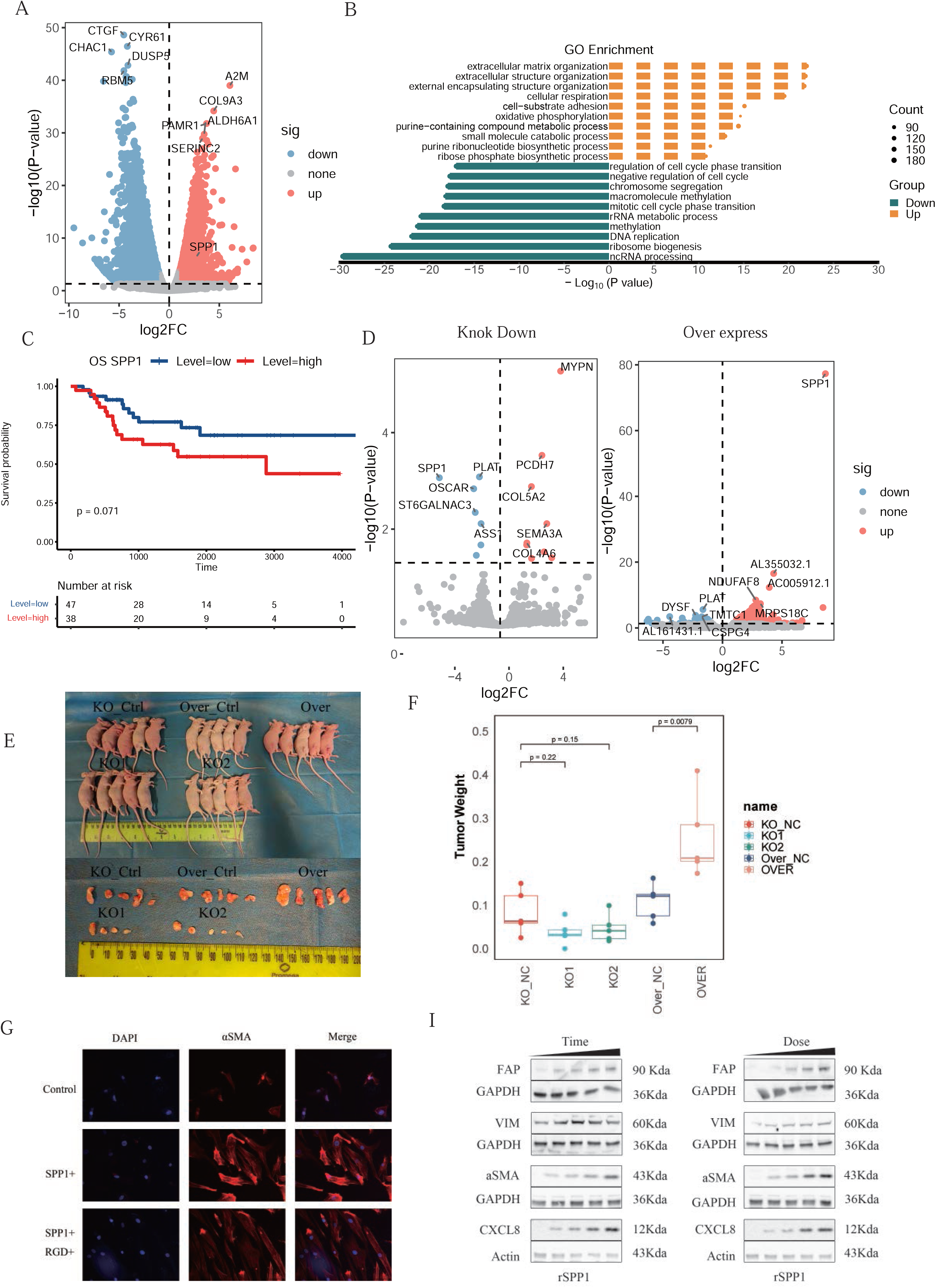
SPP1 promotes osteosarcoma proliferation and activates cancer-associated fibroblasts (CAFs). **(A)** Differentially expressed genes (DEGs) identified from bulk RNA-seq of 143B osteosarcoma cells treated with or without low-dose doxorubicin. **(B)** Gene Ontology enrichment analysis of DEGs identified from bulk RNA-seq. **(C)** Survival analysis in TARGET osteosarcoma cohort shows high SPP1 expression correlates with poor prognosis. **(D)** Bulk RNA-seq validation of SPP1 knockdown and overexpression in 143B osteosarcoma cells. **(E–F)** Xenograft models demonstrate increased tumor growth upon SPP1 overexpression. **(G)** Immunofluorescence staining of fibroblasts treated with recombinant SPP1 protein shows increased expression of α-SMA, indicative of fibroblast activation; co-treatment with RGD peptide attenuated this effect. **(H–I)** Time- and dose-dependent effects of SPP1 on fibroblast activation confirm integrin-mediated CAF remodeling.

**Figure S5.**
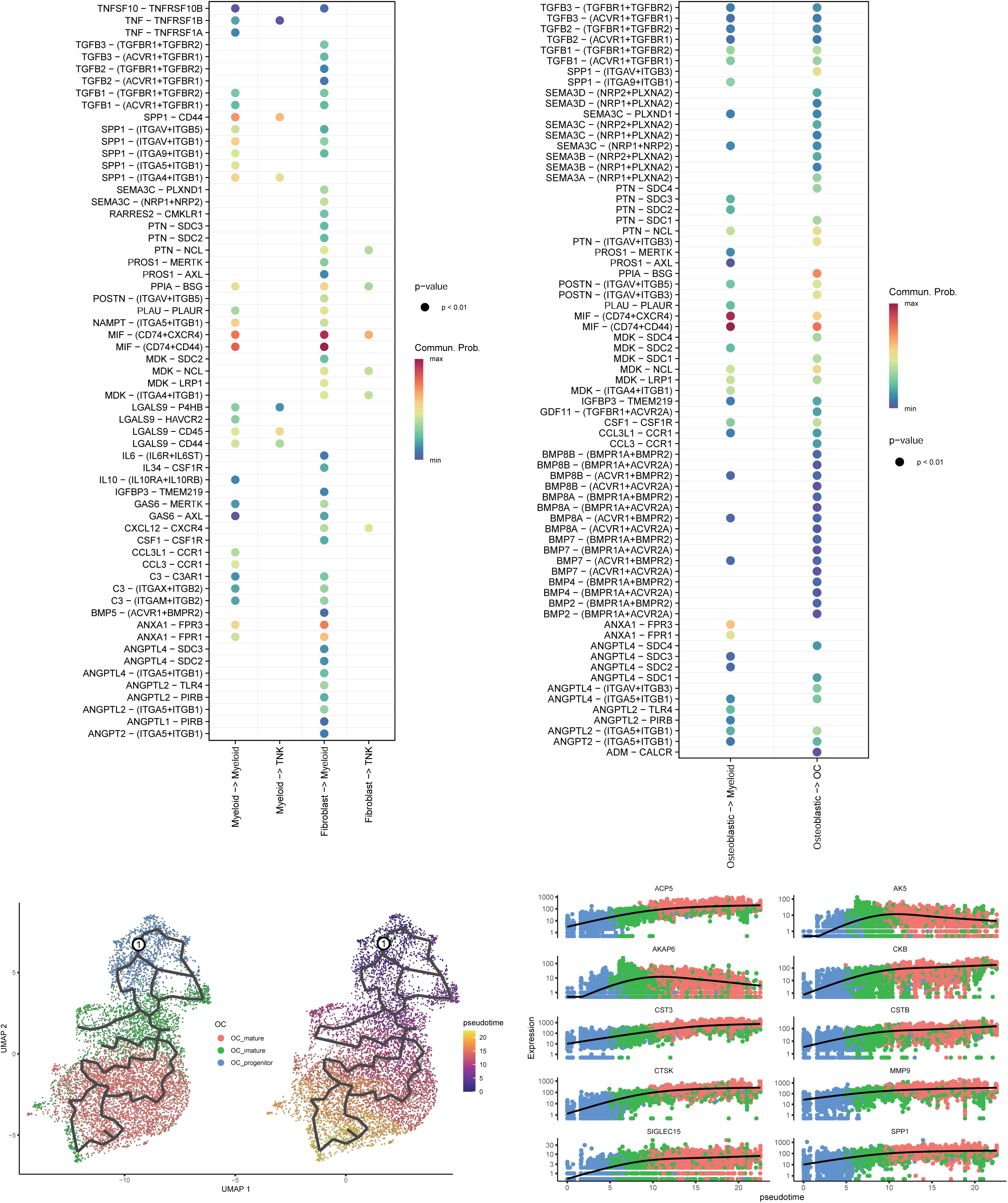
SPP1-related signaling networks in the tumor microenvironment. **(A)** CellChat analysis identified a fibroblast–myeloid–T/NK signaling axis within the osteosarcoma tumor microenvironment. **(B)** CellChat analysis identified an osteoblastic–osteoclast/myeloid signaling axis within the osteosarcoma tumor microenvironment. **(C–D)** Pseudotime trajectory of osteoclast maturation with progressive increase in ACP5, CTSK, and MMP9 expression.

Consistent with a potential role in tumor progression, high SPP1 expression was significantly associated with worse overall survival in the TARGET osteosarcoma cohort (https://ocg.cancer.gov/programs/target), underscoring its clinical relevance (Fig 5C). Functional validation experiments involving both knockdown and overexpression of SPP1 in 143B cells confirmed effective modulation at the transcript and protein levels (Fig. 5D, Fig S6B). Transcriptomic analysis of SPP1-overexpressing cells revealed enrichment of oxidative phosphorylation and respiration-related gene programs, suggesting a role in promoting mitochondrial metabolism and supporting tumor cell proliferation (Fig S6C). To assess the in vivo consequences of SPP1 upregulation, we performed subcutaneous xenograft experiments in immunodeficient mice. Tumors derived from SPP1-overexpressing 143B cells displayed significantly larger volumes compared to control tumors, while no notable differences in body weight were observed between groups, excluding systemic toxicity (Fig. 5E–F, Fig. S6D). These results demonstrate that SPP1 enhances osteosarcoma tumor growth in vivo.

Previous studies have implicated SPP1 in fibroblast activation via integrin signaling ^18, 19^. We hypothesized that tumor-derived SPP1 may contribute to tumor microenvironment remodeling by promoting cancer-associated fibroblast activation. To test this, primary human fibroblasts were treated with recombinant human SPP1 protein. Treated fibroblasts progressively acquired an activated phenotype, as evidenced by morphological changes (Fig. S6E) and increased expression of canonical CAF markers α-SMA and vimentin, as shown by immunofluorescence staining (Fig. 5G, Fig. S6F). Co-treatment with the RGD peptide, an integrin-binding motif inhibitor, abolished SPP1-induced fibroblast activation, indicating that SPP1 mediates its effects through integrin-dependent signaling (Fig. 5G, Fig. S6F). Further dose- and time-course experiments confirmed that SPP1 induces CAF activation in a concentration- and time-dependent manner (Fig. 5I, Fig. S6G).

Collectively, these results demonstrate that SPP1 functions as a dual modulator in osteosarcoma, promoting both tumor cell proliferation via metabolic reprogramming and stromal remodeling through integrin-mediated CAF activation. These effects may synergistically contribute to the establishment of an immunosuppressive tumor microenvironment.

## Discussion

In this study, we present the most comprehensive single-cell and spatial transcriptomic atlas of human osteosarcoma to date, delineating the cellular, molecular, and spatial organization of the tumor microenvironment (Fig 6). Our analysis revealed two dominant malignant phenotypes—osteoblastic and chondroblastic—that localized to distinct spatial niches shaped by microenvironmental cues. Osteoblastic-enriched regions were characterized by active osteoclastogenesis, dense fibroblast infiltration, and pronounced immunosuppressive remodeling, resembling desmoplastic subtypes previously described in osteosarcoma. In contrast, chondroblastic-dominant ecotypes exhibited reduced vascularization, hypoxia, diminished metabolic and proliferative activity, and limited immune cell infiltration. Mechanistically, we identified hypoxia-induced trans-differentiation as a key driver of chondroblastic lineage commitment, and uncovered an SPP1-mediated signaling axis through which osteoblastic tumor cells activated fibroblasts, thereby promoting stromal remodeling and immune suppression. These findings establish a spatially organized cellular ecosystem that underlies osteosarcoma heterogeneity and therapy resistance, and point to actionable targets such as the hypoxia–SOX9 axis and SPP1-mediated tumor–stroma crosstalk for potential therapeutic intervention.

**Figure 6.**
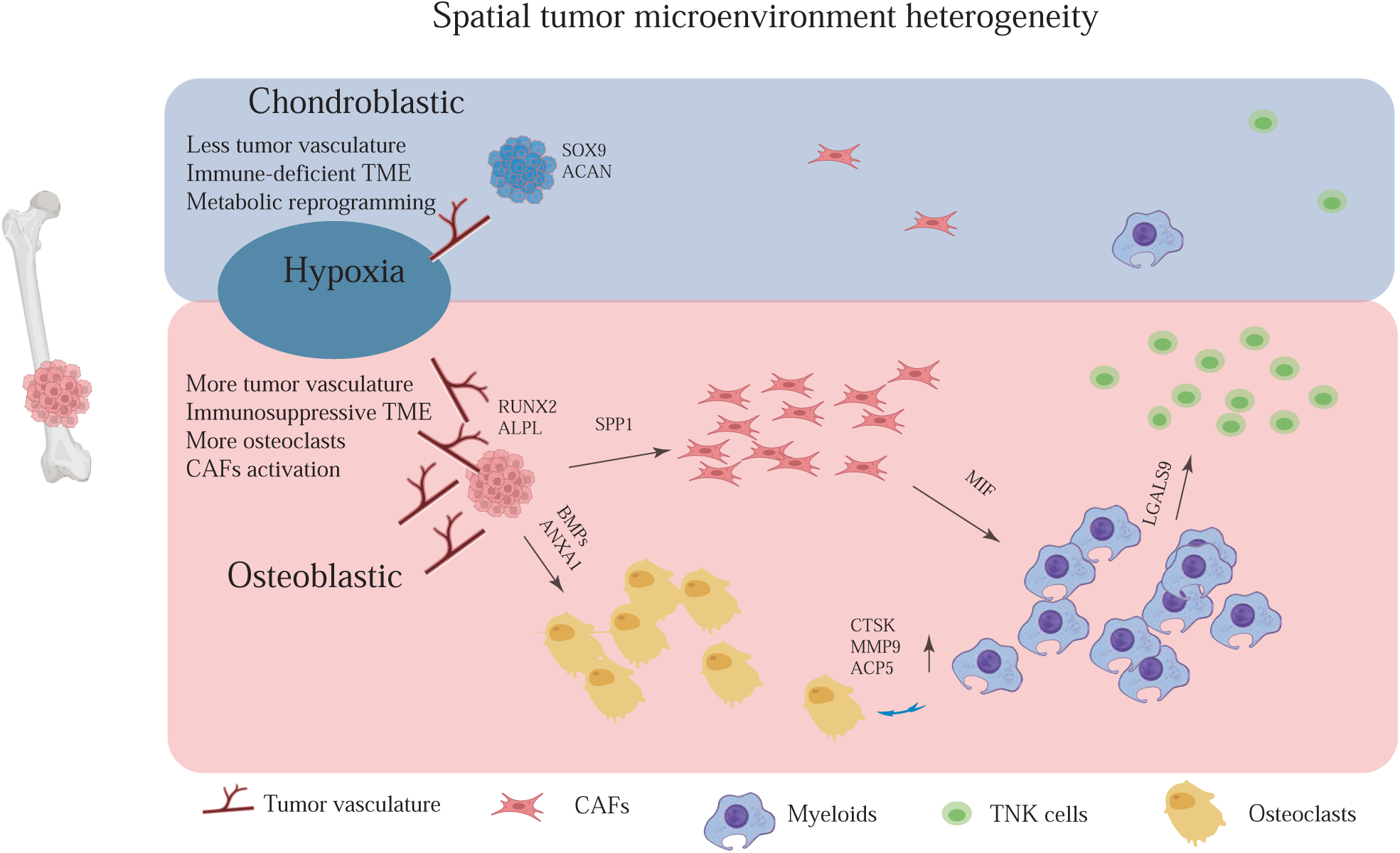
Spatially segregated osteoblastic and chondroblastic niches in osteosarcoma TME. Osteosarcoma tumor cells adopt either osteoblastic or chondroblastic differentiation trajectories shaped by distinct spatial microenvironmental contexts. Osteoblastic tumor regions are characterized by increased vascularization, metabolic activity, and infiltration of immune cells and osteoclasts. These regions express high levels of SPP1, which promotes cancer-associated fibroblast (CAF) activation through integrin signaling, leading to immunosuppressive remodeling of the TME via myeloid intermediates (MIF, ANXA1, LGALS9). Osteoclast maturation is supported by osteoblast-derived factors such as BMPs and SEMA family ligands and marked by elevated expression of CTSK, MMP9, and ACP5. In contrast, chondroblastic regions are hypoxic, poorly vascularized, and immune-excluded, with elevated HIF1A and SOX9 activity driving chondrogenic reprogramming (ACAN).

**Figure S6.**
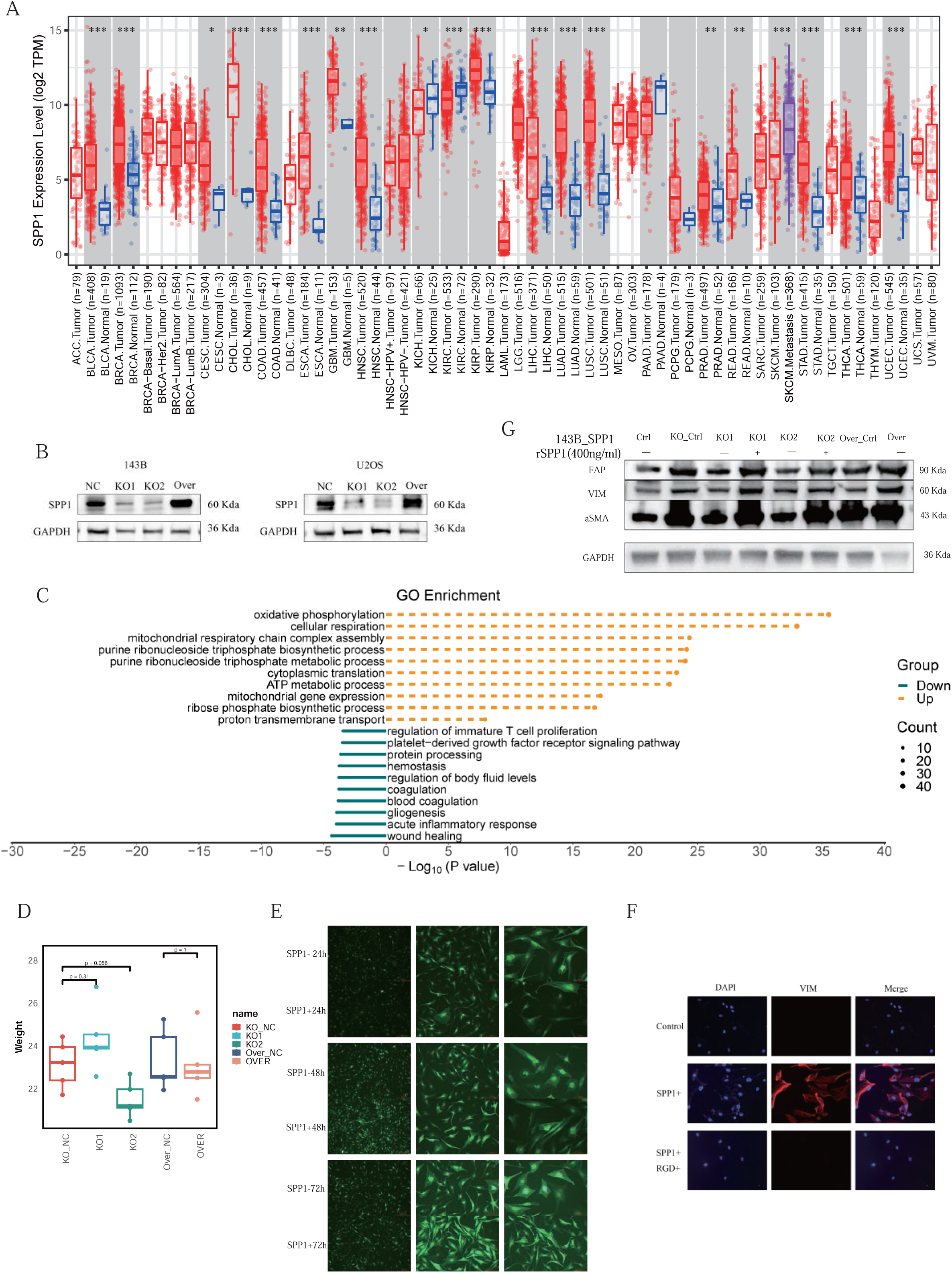
Functional validation of SPP1-mediated tumor progression and fibroblast activation. **(A)** TIMER analysis shows elevated SPP1 expression across multiple tumor types. **(B)** Western blot confirmation of SPP1 knockdown and overexpression in 143B cells. **(C)** GSVA reveals metabolic pathway enrichment in SPP1-overexpressing cells. **(D)** No significant difference in mouse body weight, confirming treatment tolerance. **(E)** Morphological changes in fibroblasts after SPP1 treatment. **(F)** Immunofluorescence staining of fibroblasts treated with recombinant SPP1 protein shows increased expression of VIM, indicative of fibroblast activation; co-treatment with RGD peptide attenuated this effect. **(G)** Genetic manipulation of SPP1 in 143B osteosarcoma cells modulates cancer-associated fibroblast activation.

Previous studies have reported lower tumor necrosis rates and worse clinical outcomes in chondroblastic osteosarcoma compared to the osteoblastic subtype ^20, 21^, suggesting distinct histology-associated behaviors. In our study, we observed the co-existence of osteoblastic and chondroblastic tumor cells within individual samples, consistent with prior reports^8^ and supportive of the concept of lineage plasticity. Importantly, previous developmental studies have demonstrated that chondrocytes can transdifferentiate into osteoblasts during endochondral ossification ^22^, suggesting that similar processes may occur in malignancy. CNV-based clonal analysis using inferCNV revealed substantial overlap in genomic alterations between osteoblastic and chondroblastic tumor subpopulations in chondroblastic OS lesions, further indicating that these lineages may arise from a shared progenitor and diverge through epigenetic, rather than genetic, mechanisms ^8^. Integrated transcriptomic and spatial analyses supported this hypothesis. While morphologically distinct, osteoblastic and chondroblastic cells displayed a transcriptional continuum and spatially organized transitions, particularly within poorly ossified invasive fronts. These patterns suggest that microenvironmental cues—most notably hypoxia—play a key role in shaping phenotypic divergence. Chondroblastic tumor cells exhibited hallmark features of metabolic suppression, including reduced mitochondrial content and proliferation, consistent with prior studies showing that malignant chondroblastic cells are less proliferative and more chemoresistant ^23^. In parallel, spatial transcriptomic data revealed that chondroblastic regions were hypovascular and hypoxic, recapitulating the physiological conditions that favor chondrogenesis. Mechanistically, we identified a hypoxia–SOX9 signaling axis as a driver of chondroblastic commitment. SOX9 is a well-established master regulator of chondrogenesis under low oxygen conditions, and its elevated expression in hypoxic tumor zones suggests a conserved regulatory programe ^24, 25^. Notably, in SP_BS7, we identified hypoxic regions at the invasive tumor front with poor ossification and high expression of both ACAN and SPP1, indicating possible trans-differentiation of chondroblastic cells toward an osteogenic phenotype. This observation aligns with Zhou et al.’s identification of transitional “Chondro_trans” cells expressing RUNX2, SPP1, and COL1A1—hallmarks of osteogenic commitment—within similar regions ^8^. Collectively, these findings suggest that the osteosarcoma differentiation landscape is highly plastic and spatially regulated, and that transitions between chondroblastic and osteoblastic states may reflect adaptive responses to dynamic microenvironmental pressures rather than rigid subclonal hierarchies.

Furthermore, our results highlight the critical role of tumor-derived secreted factors in orchestrating the stromal and immune architecture of the osteosarcoma microenvironment. Among these, SPP1 emerged as a key signaling molecule that mediates crosstalk between malignant osteoblastic cells and the surrounding stroma. SPP1 is a multifunctional glycoprotein that acts both as an extracellular matrix component and a cytokine. It has been widely implicated in promoting tumor progression, angiogenesis, metastasis, and immune modulation in a variety of cancers, including breast cancer, and pancreatic cancer ^18, 19, 26, 27^. However, its specific role in osteosarcomas has remained incompletely understood. In our study, we found that SPP1 is predominantly secreted by osteoblastic osteosarcoma cells and exerts paracrine effects on fibroblasts within the tumor stroma. Mechanistically, SPP1 engages integrin receptors on fibroblasts—likely including ITGAV, ITGB1, and CD44— to activate intracellular signaling cascades that induce a myofibroblastic transition, characterized by elevated α-SMA expression and cytoskeletal reorganization. These activated fibroblasts acquire a cancer-associated fibroblast-like phenotype, known to be involved in extracellular matrix remodeling, increased tissue stiffness, and establishment of a pro-tumorigenic niche ^28^. Importantly, these SPP1-activated fibroblasts do not merely contribute to stromal remodeling but also play a key role in shaping an immunosuppressive microenvironment. While CAFs are often associated with T cell exclusion in certain tumor types, our spatial analysis revealed a distinct pattern in osteosarcoma: CAFs co-localized not only with osteoblastic tumor cells but also with infiltrating myeloid cells and T/NK lymphocytes. Rather than forming physical barriers that exclude immune cells, CAFs in osteosarcoma appear to exert immunosuppressive influence through cellular proximity and signaling crosstalk, particularly with the myeloid compartment. CAF–myeloid interactions have been shown to foster a suppressive milieu by promoting macrophage polarization toward M2-like phenotypes ^26, 29^. In our dataset, CAF-rich regions exhibited increased enrichment of C1QA+ and SPP1+ macrophages, as well as downregulation of pro-inflammatory and interferon-related pathways ^30^. This suggests a model in which CAF-mediated immune suppression in osteosarcoma arises through immune education and tolerization, rather than simple physical exclusion. Collectively, these findings support a paradigm in which SPP1-driven CAF activation coordinates both structural and immunological remodeling of the TME, establishing a fibroblast–myeloid–T/NK signaling axis that contributes to immune dysfunction and tumor progression.

In addition to fibroblast activation, we observed strong spatial and signaling dependencies between multiple stromal and immune cell types, including endothelial cells and pericytes, myeloid cells and fibroblasts, as well as osteoclasts and osteoblastic tumor cells. This cellular architecture reflects a high degree of coordination, reminiscent of physiological bone homeostasis, where osteoblast-driven bone formation and osteoclast-mediated bone resorption are tightly coupled ^31^. Notably, osteoclasts were not only present in both primary and recurrent osteosarcoma tumors but also detected in lung metastases ^8^, indicating that osteoblastic tumor cells retain the ability to recruit and instruct myeloid cells to differentiate into osteoclasts even in ectopic, non-osseous environments. This suggests that osteosarcoma cells can hijack developmental bone remodeling programs to support invasive growth outside the bone. In line with this, our spatial transcriptomic analysis revealed marked osteoclast enrichment in osteoblastic regions, whereas chondroblastic-dominant zones exhibited significantly fewer OCs. These findings are consistent with previous studies, including Zhou et al. ^8^, which reported a paucity of osteoclasts in chondroblastic osteosarcoma. Furthermore, trajectory analysis and ligand-receptor inference suggest that osteoblastic tumor cells promote osteoclast maturation via BMPs signaling. This spatial co-localization and signaling dependency between osteoblastic cells and OCs recapitulate the osteoblast–osteoclast coupling observed in normal bone development, highlighting how osteosarcoma exploits physiological remodeling pathways to drive local invasion and tissue destruction.

Clinically, our findings offer a rationale for incorporating spatial and differentiation state information into osteosarcoma subtyping. Traditional classification based on morphology fails to capture the molecular and spatial complexity observed in our dataset. The spatial ecotypes defined here suggest that certain tumor regions may be more immunologically “cold” or chemoresistant, driven by hypoxia-induced chondroblastic transition or CAF-mediated immune suppression. Importantly, our results suggest that SPP1 and its downstream integrin signaling, as well as the hypoxia–SOX9 axis, may represent therapeutic vulnerabilities. Targeting these pathways could disrupt the stromal support and immunosuppressive architecture of aggressive OS niches. Moreover, given the elevated expression of SPP1 following chemotherapy in our models, this axis may also contribute to post-treatment relapse or resistance, further highlighting its clinical relevance.

While our cohort (n = 14 for scRNA-seq; 3 for ST) surpasses previous osteosarcoma atlases in size and resolution, larger and more diverse validation cohorts are needed to confirm our findings. Spatial proteomics or multiplexed imaging approaches are warranted to validate key ligand-receptor interactions (e.g., SPP1–CD44/integrins) at the protein level. Additionally, functional studies using osteoblast–osteoclast co-cultures may help elucidate the role of osteoclasts in tumor invasion. Finally, in vivo testing of SPP1-targeted therapies (e.g., anti-SPP1 antibodies or integrin inhibitors) in immunocompetent osteosarcoma models will be essential for translating our findings into clinical application.

In summary, this study provides an integrated single-cell and spatial framework for understanding the osteosarcoma microenvironment, identifies key cellular and signaling programs associated with distinct differentiation states, and reveals novel spatial mechanisms of immune suppression and therapy resistance. These insights lay the groundwork for developing precision therapeutic strategies that target not only malignant cells but also their supportive and immunomodulatory niches.

## MATERIALS AND METHODS

### Patients and sample collection

This study received approval from the Ethics Committee of Cancer Hospital, Chinese Academy of Medical Sciences, and Peking Union Medical College (NCC2021C-232). Fourteen patients diagnosed with osteosarcoma and treated at National Cancer Center, were recruited for participation in the study. All patients provided written informed consent.

### Single-cell library preparation and sequencing

The Single-Cell 3′ Library Kit v3 (10x Genomics) was employed for single-cell transcriptome amplification and library preparation, following procedures according to the manufacturer’s instructions. The sorted single-cell suspension was loaded onto a microfluidic chip from 10x Genomics to generate the cDNA library. Subsequently, cDNA libraries were prepared and sequenced across six lanes on an Illumina NovaSeq 6000 system (Illumina Inc., San Diego, CA, USA).

### Preprocessing of scRNA-seq data

The raw sequencing data underwent processing using CellRanger (version 7.1.0, 10X Genomics) with default parameters. The data was then mapped to the GRCh38 reference genome (Genome Reference Consortium Human Build 38), resulting in matrices of gene counts by cell barcodes. Subsequent analyses were conducted using the Seurat package ^32^ (version 4.4.0) in R (version 4.1.2, The R Foundation). Quality controls were implemented by retaining non-ribosomal genes detected in at least 0.1% of all cells, cells with a minimum of 200 features, and cells with mitochondrial fraction below 10%. Doublets were identified using the DoubletFinder package ^33^ (version 2.0.3) and subsequently removed. Normalization of raw unique molecular identifier (UMI) counts was carried out using the SCTransform function, with the number of cells set to 3,000. Harmony package ^34^ (version 1.1.0) was employed to adjust for possible batch effects arising from patient-specific expression patterns. Dimension reduction was performed through principal-component analysis (PCA) using the RunPCA function, and the optimal number of principal components (PCs) was determined using the ElbowPlot function. The same PCs were applied in cell clustering with modularity optimization using the kNN graph algorithm as input. Visualization of cell clusters was achieved using the UMAP algorithm.

### Cell type annotation

Cell types were annotated by assessing the expression of known marker genes, as detailed in previous studies, and outlined in Table. Specifically, cells expressing marker genes from at least two types of major cell types were considered undefined cells and were consequently excluded from further analysis. Cell subtypes were annotated through unsupervised clustering and examination of marker gene expression levels, as depicted in corresponding figures. Differential expression genes (DEGs) within each cell subcluster were identified using the “FindAllMarker” function with default parameters provided by Seurat. These DEGs played a crucial role in cell type annotations, where cell subclusters exhibiting similar gene expression patterns were annotated as the same cell type.

### Trajectory analysis by Monocle 3 analysis

To investigate lineage dynamics, pseudotime analysis was performed using the Monocle3 package ^35^. Preprocessed and normalized single-cell data were imported from Seurat using the SeuratWrappers interface. Cell trajectories were learned using the learn_graph function after dimensionality reduction with UMAP. Cells were ordered along the trajectory using the order_cells function, with root nodes specified based on progenitor-like cluster identity. Gene expression trends along pseudotime were visualized using plot_genes_in_pseudotime.

### Pathway analysis

Differentially expressed genes (DEGs) meeting the criteria of |logFC| > 0.5 and an adjusted *P* value < 0.05 were utilized for Gene Ontology (GO) enrichment analysis. The “compareCluster” function from the clusterProfiler package ^36^ was applied to identify significantly enriched GO terms that exhibited differences between distinct fibroblast subclusters. To evaluate the differences in pathways across distinct subsets, Gene Set Variation Analysis (GSVA) and Gene Set Enrichment Analysis (GSEA) were performed. These analyses were conducted and computed using a linear model provided by the limma package.

### Single-cell regulatory network analysis

In accordance with a standardized analysis pipeline, we employed PySCENIC ^37^ (https://github.com/aertslab/SCENIC) to examine differentially expressed transcription factors (TFs) and delve into the single-cell gene regulatory network (GRN) within distinct cell subclusters. To outline the process, we generated gene expression matrices for the cell subclusters using GENIE3, establishing initial coexpression GRNs. Subsequently, the RcisTarget package was employed to identify TF motifs within the regulon data. The AUCell package was utilized to calculate the regulon activity score for each cell. Lastly, we filtered regulons with a correlation coefficient exceeding 0.3 with at least one other regulon, opting for those activated in a minimum of 30% of the cell subclusters for subsequent visualization.

### Differentiation of tumor cells and nonmalignant cells based on inferCNV

To distinguish tumor cells from nonmalignant cells, we employed the inferCNV R package (https://github.com/broadinstitute/inferCNV) to estimate the initial copy number variation (CNV) signal for each of the 60 regions using default parameters. The CNV signal was computed as the quadratic sum of the CNV region, and cells with a CNV signal surpassing 0.04 were designated as potential tumor cells in the context of OS (osteosarcoma).

### Gene set variation analysis

Pathway analyses were predominantly carried out on specific GO/Hallmark pathways obtained from the Molecular Signature Database (MSigDB version 6.2). To gauge pathway activity at the individual cell level, we utilized gene set variation analysis (GSVA) with default settings, implemented through the GSVA package (version 1.34.0).

### Gene set signature scoring

To quantitatively represent each cell subtype, we determined a subtype signature score, defined as the arithmetic mean of the expression levels of signature genes sourced from the specified data slot (e.g., SCT data slot). This methodology facilitates a standardized and comparative analysis of cell subtypes across various conditions and studies. Signature scores were computed using AUCell with default parameters ^38^. The genes employed for gene set signature scoring are detailed in Supplementary Table 2.

### Cell-cell interaction analysis using Cellchat

To investigate the cell-cell interaction network between different cell types, we used CellChat v2 (https://github.com/jinworks/CellChat) in combination with the CellPhoneDB receptor-ligand database. The analysis was performed following established protocols to identify potential receptor-ligand interactions between different cell subtypes.

First, we obtained a list of receptors and ligands expressed in the cells. For each cluster, only those receptors and ligands that were expressed in more than 10% of the cells within that specific cluster were considered for the analysis. This filtering ensured that the interactions were biologically relevant and not driven by rare or stochastic gene expression. Cell-cell interactions were then inferred using the CellChat package, which relies on the CellPhoneDB database to identify significant receptor-ligand pairs across the cell populations. The interactions were further filtered based on the expression levels and interaction scores derived from the database.

### Tumor ecotype analysis using deconvolution

CIBERSORTx (https://cibersortx.stanford.edu/) ^39^ were used to deconvolute predicted cell-fractions from a number of bulk transcript profiling datasets. This algorithm employs a support vector machine (SVM) regression approach to estimate the relative proportions of various cell types based on gene expression signatures. We applied CIBERSORTx to predict the cellular composition in osteosarcoma samples by comparing the bulk transcriptomic data with predefined cell-type signature matrices. The deconvolution process generates estimates of cell-type fractions, and permutation testing (1000 iterations) was used to assess the robustness of these predictions. The results provided insights into the tumor microenvironment and highlighted variations in cellular composition across different tumor samples.

### Spatial transcriptomic sequencing

We performed 10x Genomics Visium spatial transcriptomic sequencing on osteosarcoma samples having high RNA quality and recognizable histopathological status, using the methods as described ^40^. Sequencing was conducted using the Illumina NovaSeq 6000 platform (Illumina). Raw sequencing data were processed through the Spaceranger workflow (version 2.0.0) for alignment and quantification. The resulting count matrices were then loaded into the Seurat ^32^ and RCTD ^41^ packages for subsequent data filtering, normalization, dimensional reduction, and visualization.

For spatial tissue deconvolution, we used annotated single-cell RNA sequencing (scRNAseq) data from 14 osteosarcoma patients, initially processed using Seurat and analyzed through cell2location ^42^ to resolve the spatial organization of cell types across the tissue sections. Further analysis was performed using Harmony ^43^ for batch correction, where gene expression data were integrated to perform clustering and identify 12 distinct cell clusters. The relationship between niche and cell type co-localization was then assessed using correlation analysis. Additionally, the Misty algorithm was applied to evaluate the importance of different cell types within the osteosarcoma microenvironment.

### Western blot

To perform a Western blot assay, total protein was extracted from cell lysate. The protein level was determined using a BCA kit or stained in gel and scanned. The proteins were then subjected to SDS-PAGE, transferred to a PVDF membrane, and probed with various antibodies from Abcam or for Proteintech specific proteins like OPN, Vinculin, SOX9, HIF1A, CD16, etc. After incubation with primary and secondary antibodies, the membrane was visualized using a chemiluminescent substrate in an imager.

### Immunofluorescence analysis

Tissue sections were deparaffinized, rehydrated, and subjected to antigen retrieval. After blocking non-specific binding with a suitable blocking agent, sections were incubated with primary antibodies overnight at 4°C. Following washing, fluorescently-labeled secondary antibodies were applied. Nuclei were stained with DAPI. Slides were mounted and images captured using a fluorescence microscope. Controls included sections incubated without primary antibodies to assess non-specific binding of secondary antibodies.

### Immunohistochemistry analysis

Tissue sections were deparaffinized and rehydrated through graded alcohols to water. Antigen retrieval was performed using citrate buffer in a microwave. Endogenous peroxidase activity was quenched with hydrogen peroxide. After blocking with normal serum, sections were incubated with primary antibodies overnight at 4°C. Biotinylated secondary antibodies were applied, followed by amplification with an avidin-biotin complex. Color development was achieved with DAB substrate. Sections were counterstained with hematoxylin, dehydrated, and mounted. Controls omitted the primary antibody.

### Animal models

Female Balb/c-nu nude mice (4–5 weeks old, ∼18 g, SPF (Beijing) Biotechnology Co., Ltd.) were housed under standard conditions (22 ± 2°C, 50 ± 10% humidity) with ethical approval from National Cancer Center. The 143B osteosarcoma cells (control groups: KO_Ctrl, Over_Ctrl; SPP1-knockdown groups: KO1, KO2; SPP1-overexpression group: Over) were cultured in DMEM with 10% FBS, harvested during logarithmic growth, trypsinized, washed twice with PBS, and resuspended to 4 × 10⁶ cells/100 μL. Viable cells (>95% by trypan blue) were subcutaneously injected (100 μL) into the right inguinal region of ethanol-disinfected mice using a 29G insulin syringe, with pre-injection plunger retraction to avoid vascular puncture. Tumor dimensions (L, W) were measured every 4 days for 31 days, and volume calculated as V = 0.5 × L × W². Mice were euthanized by CO₂ asphyxiation at humane endpoints, and tumors were excised, weighed, fixed in 4% PFA, and processed for histology.

### Statistical Analysis

The in the study were conducted using R 4.3.0 and Python 3.9.0. Specific statistical details and methods are described in the figure legends, main text, or methods section. P-values were calculated using a two-sided, unpaired Wilcoxon rank-sum test. For error representation, either standard error of the mean (S.E.M.) or standard deviation (S.D.) was used, based on a minimum of three independent experiments.

